# NetREx – Network-based Rice Expression Analysis Server for Abiotic Stress Conditions

**DOI:** 10.1101/2021.12.24.474096

**Authors:** Sanchari Sircar, Mayank Musaddi, Nita Parekh

## Abstract

Recent focus on transcriptomic studies in food crops like rice, wheat and maize provide new opportunities to address issues related to agriculture and climate change. Re-analysis of such data available in public domain supplemented with annotations across molecular hierarchy can be of immense help to the plant research community, particularly co-expression networks representing transcriptionally coordinated genes that are often part of the same biological process. With this objective we have developed NetREx, a Network based Rice Expression Analysis Server, that hosts ranked co-expression networks of *Oryza sativa* using publicly available mRNA-seq data across uniform experimental conditions. It provides a range of interactable data viewers and modules for analysing user queried genes across different stress conditions (drought, flood, cold and osmosis) and hormonal treatments (abscisic and jasmonic acid) and tissues (root and shoot). Subnetworks of user-defined genes can be queried in preconstructed tissue-specific networks, allowing users to view the fold-change, module memberships, gene annotations and analysis of their neighborhood genes and associated pathways. The webserver also allows querying of orthologous genes from Arabidopsis, wheat, maize, barley, and sorghum. Here we demonstrate that NetREx can be used to identify novel candidate genes and tissue-specific interactions under stress conditions and can aid in the analysis and understanding of complex phenotypes linked to stress response in rice. Available at: https://bioinf.iiit.ac.in/netrex/index.html

## 1. Introduction

Entities in a biological system seldom work in isolation. There is a coordinated association between them at every level of the molecular hierarchy. These associations are context-specific and involve making and breaking of links among the entities depending upon the developmental stage, tissue, and environmental conditions. In field conditions, plants are generally exposed to multiple stresses. Thus, finding processes that are common or unique across various stress conditions can provide novel insights towards multi-stress tolerant crops. One of the first studies on multi-stress expression profiling in rice (using rice cDNA microarray) was conducted by Rabbani *et al* (1) with the objective to identify genes induced by cold, drought, high-salinity, and abscisic acid (ABA) treatments. Although this early study was on a limited set of probes, it demonstrated common set of genes differentially expressed across various environmental conditions. This paved the way for numerous studies on various crops: expression profiling in potato in response to cold, heat and salt stress (2), effect of ABA, drought and salinity stress on pathogen defense in tomato (3), transcriptome responses during single and combination of stresses in Arabidopsis (4), expression profiling of chickpea genes in response to salinity, drought and cold stress (5), etc.

Transcriptomic resources have made major contributions in plant functional genomics. For example, RiceXPro hosts rice transcriptomic data across various conditions, *viz*., different plant hormone treatments, cell and tissue types, and field/developmental stages (6). The database RiceSRTFDB provides expression information for transcription factors (TFs) in rice under drought and salinity stress and various developmental stages (7). More recent resources include ePlant webserver that provides an integrated view of the genome, proteome, interactome, transcriptome (both from microarray and RNA-seq data) and 3D molecular structure data in Arabidopsis (8), Rice Expression Database (RED) (9), TENOR database, a collection of large-scale mRNA sequencing (mRNA-seq) data obtained from rice under a wide variety of conditions (10), etc. For researchers, such large-scale resources provide further opportunities to re-analyse the data with new hypotheses.

A transcriptomic experiment (microarray or RNA-Seq) typically detects thousands of genes differentially expressed with respect to control conditions. Functional analysis of such a large set of genes is a daunting task. A popular approach is to construct co-expression networks based on the similarity of gene expression profiles (11). Here, nodes represent genes and edges represent the strength of correlations between them. Highly correlated genes can then be grouped into functional modules, reducing the dimensionality of data. Another approach is to convert raw correlation values (Pearson or Spearman) to highest reciprocal ranks (HRRs) or mutual ranks (MRs) (12,13). Several studies have shown that the biological relevance of ranked-based methods are more significant and robust than those constructed using raw Pearson correlation coefficients (PCCs) because they intuitively consider PCCs of the neighbourhood of gene pairs as well (13–15). Moreover, these methods circumvent the problem of hard-thresholding of PCCs and allow retaining of weak but significant co-expression relationships between genes on the basis of ranks (16–19).

For the plant community, various functional resources are available. At the global-scale, *condition-independent* networks constructed across a large number of datasets from varying experimental conditions, platforms, tissues and developmental stages are available, e.g., AraNet (20), RiceNet (21), MaizeNet (22). Although these allow one to derive associations from a large sample size, the merging of different experimental conditions might incorporate complexities in the network difficult to account for. Nevertheless, these resources have been useful in functional annotation of unknown genes and prioritization of candidate genes (20–22). On the other hand, *condition-dependent* networks, derived from smaller datasets representing specific conditions, provide opportunities to query context-specific associations (23,24). For the construction of this resource we have considered publicly available mRNA-seq data from TENOR database (10) that hosts genome-wide time-course transcriptome study of rice shoots and roots under a wide variety of conditions, *viz*., environmental stresses (e.g., drought, cold, flood, salinity), nutrient deficiencies and excesses (e.g., phosphate), heavy metal toxicity stress (e.g., cadmium) and plant hormone treatments (abscisic acid (ABA) and jasmonic acid (JA)) have been used. All the experiments have been performed using a single mRNA-Seq platform under standardized laboratory conditions and from the same rice cultivar *Oryza sativa L*. (cv. Nipponbare), making it possible to compare gene expressions across different stress conditions. In NetREx, data for four stress conditions (drought, cold, flood and osmotic stress) and two hormonal treatments (ABA and JA) from seedlings (10 days after germination) have been considered for network construction and analysis. First, we construct a global rank-based stress network across four stress conditions and two hormonal treatments, separately for root and shoot tissues. Stress-specific networks are then derived from these global networks using differentially expressed genes. These networks along with functional information are hosted in the database.

## 2. Materials and Methods

### 2.1. Datasets

Nucleotide sequences corresponding to mRNA-seq data from rice seedlings (*Oryza sativa L. ssp. japonica cv. Nipponbare*) under various conditions are obtained from DDBJ Sequence Read Archive (DRA000959). These are 76 bp single-read sequences obtained from Illumina GAIIx instrument under uniform library conditions (10). The details of the 7 datasets are summarized in **Table 1**.

**Table 1:** Summary of mRNA-Seq data from DDBJ-SRA considered for multi-stress analysis.

| Conditions | Treatment | Time-points (No. of biological replicates) |  | No. of Samples |
| --- | --- | --- | --- | --- |
|  |  | Root | Shoot |  |
| **Cold** | 4°C | 0h (3), 1h (2), 3h (2), 6h (2),<br>12h (2), 1d (2) | 0h (3), 1h (2), 3h (2), 6h (2),<br>12h (2), 1d (2) | Root:13<br>Shoot:13 |
| **Drought** | Grown without medium | 0h (2), 1h (2), 3h (2), 6h (2),<br>12h (2), 1d (2) | 0h (2), 1h (2), 3h (3), 6h (2),<br>12h (2), 1d (2) | Root:12<br>Shoot:13 |
| **Osmotic** | 0.6 M Mannitol | 0h (3), 1h (2), 3h (2), 6h (2),<br>12h (2) | 0h (3), 1h (2), 3h (2), 6h (2),<br>12h (2) | Root:11<br>Shoot:11 |
| **Flood** | Completely submerged in medium | 0h (3), 1h (2), 3h (2), 6h (2),<br>12h (2), 1d (2), 3d (2) | 0h (3), 1h (2), 3h (2), 6h (2),<br>12h (2), 1d (3), 3d (2) | Root:15<br>Shoot:16 |
| **ABA** | 100 µM | 0h (2), 1h (2), 3h (2), 6h (2),<br>12h (2), 1d (2) | 0h (2), 1h (2), 3h (2), 6h (2),<br>12h (2), 1d (2) | Root:12<br>Shoot:12 |
| **JA** | 100 μM | 0h (2), 1h (2), 3h (2), 6h (2),<br>12h (2), 1d (2) | 0h (2), 1h (2), 3h (2), 6h (2),<br>12h (2), 1d (2) | Root:12<br>Shoot:12 |
| **No**<br>**Treatment**<br>**(NT)** | - | 0h (2), 1h (2), 3h (2), 6h (2),<br>12h (2), 1d (2), 3d (2), 4d (2),<br>5d (2), 10d (2) | 0h (2), 1h (2), 3h (2), 6h (2),<br>12h (2), 1d (2), 3d (2), 4d<br>(2), 5d (2), 10d (2) | Root:20<br>Shoot:20 |
| **Total** |  |  |  | **Root:95**<br>**Shoot:97** |

### 2.2. Pre-processing

Raw reads are processed for quality control. Adaptor sequences and low-quality bases (< Q15) at 5′ and 3′ ends of the reads are trimmed using Cutadapt (ver. 1.15) (25). After trimming, reads < 20 bp are discarded as these may lead to non-specific alignment to reference genome. For each sample, reads are aligned to the rice reference genome (Os-Nipponbare-Reference-IRGSP-1.0) using graph-based alignment program, HISAT2 (ver. 2.1.0) (26) and gene annotations are taken from RAP-DB database (version: 2017-04-14) (27). For each sample total base pairs before and after filtering and percentage of reads aligned to the genome are given in **Table S1-S7**. The percentage of reads mapped to the reference is a global indicator of sequencing accuracy as well as the presence of contamination in the samples (28).

### 2.3. Estimating read counts and differential gene expression analysis

Raw read counts for a gene are a measure of its expression. Here we use ‘featureCounts’ tool from SubRead package (1.6.0) (29) and gene annotation file from RAP-DB to compute gene expression. Percentage of reads assigned to the genes is given in the last column in **Table S1-S7**. Differentially expressed genes (DEGs) are determined in a tissue-specific manner for every time-point at 2-fold change and *p-*value < 0.05 using DeSeq2 (30). Genes that are differentially expressed in at least 2 time-points for a given stress condition are given in **Table 2**.

**Table 2:** Number of differentially expressed genes (DEGs) across various conditions.

| Tissue\Stress | DEGs in at least 2 time-points |  |  |  |  |  |  | Total DEGs – NT | Total DEGs used for network construction |
| --- | --- | --- | --- | --- | --- | --- | --- | --- | --- |
|  | Drought | Cold | Osmotic | Flood | ABA | JA | NT |  |  |
| **Root** | 3817 | 6327 | 8040 | 3651 | 9340 | 8835 | 1127 | 16812 | 13695 |
| **Shoot** | 10576 | 4540 | 6267 | 6955 | 8898 | 6531 | 1116 | 15804 | 13717 |

Stress/treatment-specific DEGs overlapping with the developmental time-points (NT) are filtered out as these basically account for diurnal/developmental changes. Final count matrix across various stress/hormone treatments is constructed considering genes having raw read counts ≥ 5 in at least 50% of the total samples across time-points. In RNA-seq data, most strongly expressed genes show large variations across samples compared to those with lower expression profiles (heteroscedasticity). On the other hand, most common methods of multi-dimensional data analysis like clustering work best with homoscedastic data (variance is independent of mean). To achieve this approximate homoscedasticity, the combined count matrix is normalized using variance stabilizing transformation (VST) in DeSeq2 to obtain a relatively flat trend of variance as a function of mean (31). The DEGs are then considered for rank-based network construction (details given in Sec. 2.4). The **Table 2** below indicates that for certain stress conditions, e.g., drought and flood stress, a large number of DEGs are observed in shoot tissue compared to root tissue, indicating that major physiological changes like growth in the green tissues and plant height get impacted by these conditions. Also, exogenous and direct application of hormones like ABA and JA induce drastic changes on the plant at the molecular level with large number of DEGs getting activated or repressed.

### 2.4. HRR Network Construction

A network-based approach is used to capture the associations between genes that are up- or down-regulated under various stress conditions. For this, Pearson correlation coefficients (PCCs) are computed between differentially expressed genes under various stress conditions (**Table 2**), for root and shoot tissues separately. Positively correlated genes with *p*-value < 0.05 are considered for the construction of Highest Reciprocal Rank (HRR)-based co-expression network proposed by Mutwil *et al* (12), both for root and shoot tissues. The HRR score between genes *A* and *B* is given by:

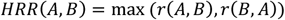

where *r*(*A, B*) is correlation rank of gene *B* in gene *A*’s co-expression list and *r*(*B,A*) is correlation rank of gene *A* in gene *B*’s co-expression list. For this study, the root and shoot networks are constructed with HRR values ≤ 100 (i.e., only top 100 neighbours for each gene are considered) and is termed as ‘HRR-100’ network. Corresponding stress-specific sub-networks are derived from the HRR-100 network for each of the tissues using respective stress-specific DEGs in **Table 2**.

Number of DEGs across various conditions in the correlation network and HRR-100 network are given in **Table 3** and **Table 4** respectively. It maybe be noted that the major advantage of considering HRR-100 network over PCC network is a significant reduction in the number of edges. Moreover, while ranking the genes, this process intuitively considers the vicinity of the network-neighbors as well which has a considerable impact on the biological relevance of the networks (14).

**Table 3:** Number of DEGs across various conditions in Pearson Correlation Coefficient (PCC) network.

| Tissue \ Stress |  | PCC network (with PCC > 0 and p-value < 0.05) |  |  |  |  |  |  |
| --- | --- | --- | --- | --- | --- | --- | --- | --- |
|  |  | Total Network | Drought | Cold | Osmotic | Flood | ABA | JA |
| Root | Nodes | 13695 | 2970 | 5005 | 6311 | 2347 | 7460 | 7565 |
|  | Edges | 2,62,69,920 | 16,86,453 | 36,37,333 | 62,86,236 | 8,95,590 | 83,63,871 | 96,29,038 |
| Shoot | Nodes | 13717 | 8809 | 3688 | 5280 | 5791 | 7546 | 5523 |
|  | Edges | 2,91,80,730 | 1,44,25,817 | 24,30,142 | 49,56,792 | 57,93,144 | 95,82,721 | 53,20,944 |

**Table 4:** Number of DEGs across various conditions in HRR-100 network.

| Tissue \ Stress |  | HRR-100 |  |  |  |  |  |  |
| --- | --- | --- | --- | --- | --- | --- | --- | --- |
|  |  | Total Network | Drought | Cold | Osmotic | Flood | ABA | JA |
| Root | Nodes | 13695 | 2970 | 5005 | 6311 | 2347 | 7460 | 7565 |
|  | Edges | 21,38,990 | 1,30,207 | 3,52,715 | 3,53,129 | 63,630 | 4,63,175 | 4,66,528 |
| Shoot | Nodes | 13717 | 8809 | 3688 | 5280 | 5791 | 7546 | 5523 |
|  | Edges | 22,72,758 | 6,57,747 | 1,78,198 | 3,03,953 | 3,35,654 | 5,00,651 | 3,21,672 |

### 2.5. Network Clustering with WGCNA

Co-expressed gene clusters often point at coordinated biological processes and help in reducing the dimensionality of the data. For this purpose, ‘signed’ co-expression network is constructed with 13695 and 13717 DEGs from root and shoot tissues respectively. The unsigned networks use absolute value of correlations, 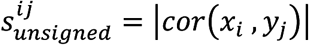, and are unable to distinguish between gene activation (s^ij^_unsigned_ = 1) and gene repression (s^ij^_unsigned_ = 1), leading to loss of biological information (32). Hence, here we construct a signed co-expression network, considering the ‘sign’ of correlation between expression profiles of genes and the similarity measure in this case is defined as:

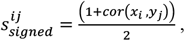

where *x*_*i*_ and *x*_*j*_ are the expression profiles of genes *i* and *j* across the samples. Here, s^ij^_signed_ = 1 corresponds to positive correlation, s^ij^_signed_ = 0, negative correlation and s^ij^_signed_ = 0.5, no correlation, thereby distinguishing between positively and negatively correlated genes.

The function *block-wiseModules* in WGCNA R package is used for hierarchical clustering of genes using Dynamic Tree Cut approach (33) with maximum block size = 14000, minimum module size = 50, “cut height” = 0.995 and “deep split” = 2. The tissue-specific network parameters are given in **Table 5**. Here, for a weighted network, “**β”** is the soft thresholding power to which co-expression similarity is raised to calculate adjacency, thereby emphasizing high correlations at the expense of low correlations. And “**k**” refers to the connectivity or sum of the connection strengths with other genes in the network.

**Table 5:**
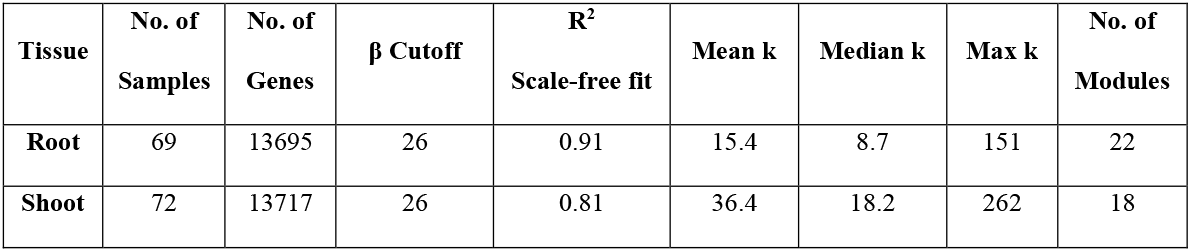
Parameters used for the construction of signed, weighted gene co-expression network using WGCNA R package is summarized.

The co-expressed modules (22 for root and 18 for shoot) are submitted for over-representation analysis in GO consortium using PANTHER classification system (v14.0) (34) for *Oryza sativa*. Results with Fischer’s Exact Test and Bonferroni correction for multiple testing (*p*-value < 0.05) are retrieved for each module. The weighted topological overlap matrix in WGCNA allows us to compute various degree centrality measures which can be useful in screening important genes. Specifically, we computed k_Total_, the weighted connectivity of a gene in the whole network and k_IM_, the within-module degree or intramodular connectivity of a gene in respective co-expressed module.

## 3. Server Implementation and GUI

Visualization and interpretation of large-scale datasets to study the emergent properties of complex biological processes with functional annotations across the molecular hierarchy has become an indispensable task for the scientific community. With this objective, we have developed NetREx, the web-based network querying and visualization resource (**Figure 1**). Using NetREx one can view relationships between query genes and analyze them based on different supported visualizations like the network viewer, the network neighborhood viewer and the expression viewer. It also provides the option to browse through the complete database based on tissue, modules, stress conditions/ hormone treatment and KEGG pathways. It is built using Sigma JS(3), a javascript library dedicated to graph drawing. The backend has been built on Express JS that communicates between the user and the JSON database and allows handling of all logical computations like identifying the query, fetching and filtering of data from the JSON database that stores the complete graph, and extract relationship between queried genes for chosen tissue and stress/treatment conditions. Clustergrammer JS is a front end javascript library for building heatmap plots in the Expression Viewer and provides interactive visualization features such as sorting and zooming (35). To make the querying efficient and fast, the gene node’s position, size, shape and color are pre-calculated and stored in JSON files using the Gephi software (34).

**Figure 1.**
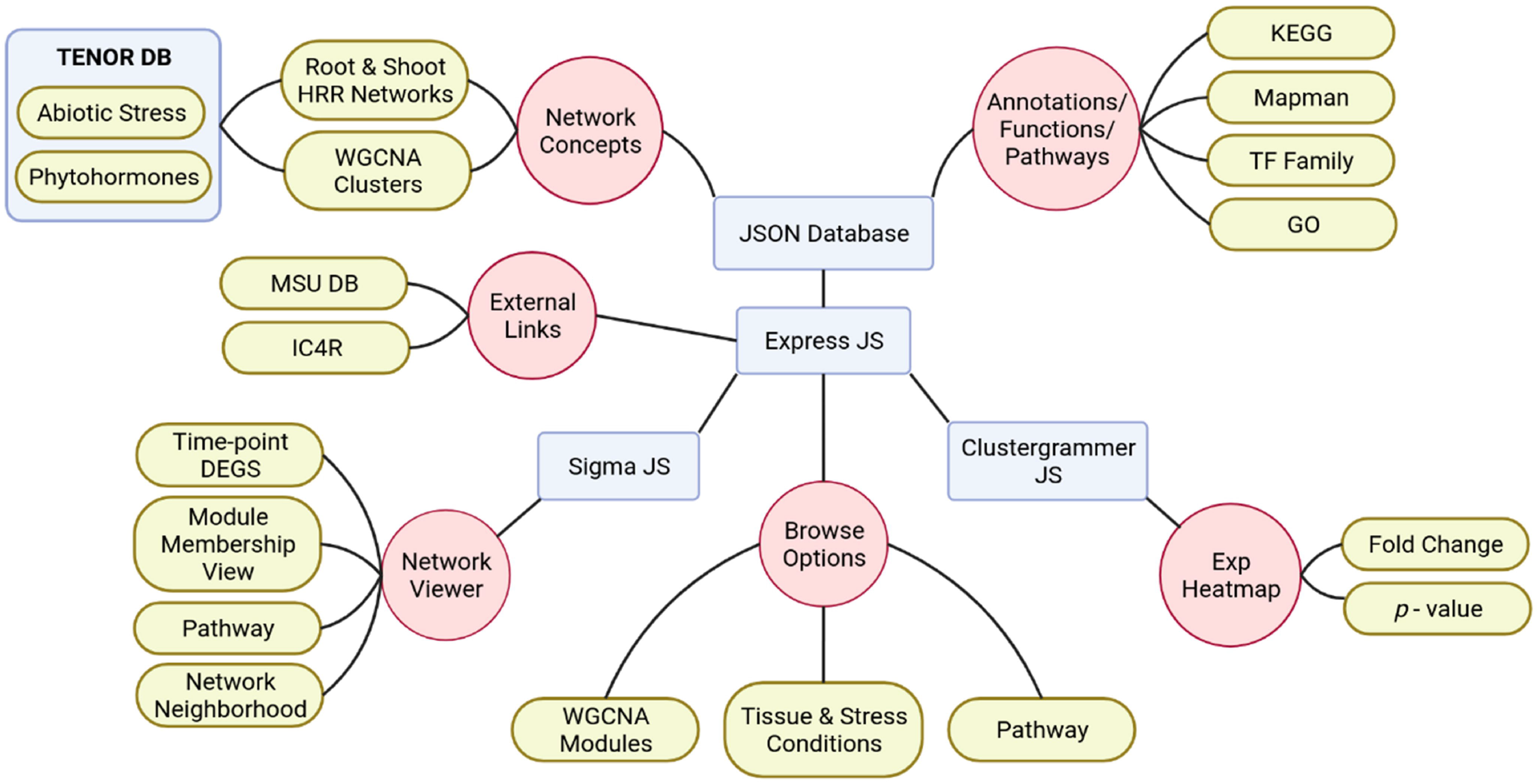
Schematic representation of NetREx architecture, modules and visualizations.

### 3.1. Modules and Views

The interface of NetREx is kept simple for the ease of use for the biologists. A comma-separated list of gene IDs (in RAP-DB format) can be uploaded or manually pasted. A user may navigate NetREx either by querying a chosen set of genes or by browsing through various modules. Below we discuss in detail various functionalities of NetREx.

#### Querying NetREx

A user can submit up to a maximum of 300 genes and query their expression profiles across any of the four abiotic stresses, *viz*., drought, cold, osmosis and flood, and two phytohormone treatments, abscisic acid and jasmonic acid, for root or shoot tissues. In the dropdown menu, the user may provide rice RAPDB IDs. Alternatively, the user may also query using Ensembl Stable IDs for Arabidopsis, wheat (*Triticum aestivum*), maize (*Zea mays*), barley (*Hordeum vulgare*) or sorghum (*Sorghum bicolor*), and the genes will be mapped to the corresponding rice orthologs based on the Ensembl Plants database. This allows the user to investigate involvement of genes in other crops under abiotic stress and phytohormone treatments through the interaction networks of their orthologs in rice.

#### Expression Viewer

Early and late stress-responsive genes are known to have distinct roles in stress-response. At the same time, genes that are ubiquitously differentially expressed across all the stages of plant may have some essential roles (34). Earlier studies have shown tissue-specific roles of stress-responsive genes indicating that divergence in the expression patterns of differentially expressed genes is an important indicator of their functions. Particularly, for uncharacterized genes, stage and tissue-specific expression profiles can give important cues regarding their functions (36). The user can analyze such stress, tissue and time-point specific information through heat maps provided in the Expression Viewer module. For the user provided gene set, differential expression of genes can be observed based on fold-change and *p*-values at various time-points for the chosen stress and tissue. The user is also provided with the option to sort the ordering of genes or time points in the heatmap based on the expression value, thus allowing to observe general trends across them.

#### Network Viewer

This module displays connectivity between query genes in different stress/treatment conditions and tissue type at different time-points. The panel on the right provides various functionalities and colouring schemes to assess the relevance of the nodes. The size of the node is drawn proportional to its connectivity, i.e., larger the node, higher is its connectivity. Also, two different shapes are used for the nodes, ‘circle’ for genes and ‘triangle’ for transcription factors. In the “default view” a gradient colouring scheme based on the degree of nodes is displayed. Thus, darker the node colour, higher is its degree, i.e., connectivity. Owing to the dense nature of the network, the arrangement of nodes in the area plays an important role in how the network is inferred. Thus, to make highly connected gene nodes appear together and unrelated nodes drift apart, a force directed algorithm, ForceAtlas2 has been used. Upon hovering over any node in the network, it along with its first neighbours get highlighted. The nodes may also be coloured based on its expression status: ‘red’ or ‘blue’ depending on whether the gene is up- or down-regulated at a chosen time point. Thus, by comparing across different time-points, the user can analyse which genes go up or down as a function of time. This functionality helps in identifying ‘early’ or ‘late’ responsive genes. This feature can also be used to compare differential expression of genes across tissues, or across different stress/treatment conditions. Another visualization option provided in NetREx is to colour nodes according to their module membership in WGCNA clusters. If majority of query genes are part of the same co-expressed module, then it is highly likely that they represent the same biological process and based on their up or down regulation, we can know whether the process is activated or repressed.

The user may also highlight the genes based on KEGG pathway categories provided in the dropdown menu. Further, to aid in the identification of genes responsive to multiple stress conditions, the user can load other stress conditions from the pull-down menu and view results in a new tab. Additional information, based on “Node” and “Edge” attributes, is provided in a tabular format by clicking on the “Show/Hide Table” button. In the “Node” attributes table, gene attributes such as gene IDs (RAPDB and MSU), transcription factor (TF) annotations, module membership, gene descriptions and link to IC4R Rice Expression Database (RED) (9) are provided. Access to IC4R allows the users to compare gene expression profiles across a larger set of RNA-seq experiments (24 projects) across various growth stages, tissues, and conditions. Gene functional annotations such as GO terms (based on RAPDB annotations) and pathway annotations from KEGG databases (37) and MapMan (38) are also provided. The fold-change and *p*-values for each gene for every time-point for the chosen tissue and stress/treatment condition is given. In the “Edge Attributes” table, Pearson Correlation Coefficients (PCC) and HRR ranks for each interacting pair are given for the selected tissue-specific network. On the right panel, the user can view up to 100 top neighbours (default=50) of the query genes based on the k_Total_ (connectivity in the whole tissue-specific co-expression network). The query genes are depicted as nodes encircled with green border.

#### Browsing NetREx

##### By Modules

Using WGCNA R package 22 co-expressed gene modules for root and 18 for shoot HRR networks were identified (**Table 5**). These gene modules can be accessed by clicking “Module-wise” under the “Browse” menu and choosing the tissue and module name from drop down menu. On this page, top 100 highly connected genes based on their within-module connectivity can viewed on a graded colour scale, based on the “colour name” of the module. The top 100 genes of the module are also listed in tabular format along with their node and edge attributes, functional annotation (from GO, MapMan and KEGG) and GO enrichment. For a given module in a tissue-specific network, these genes represent the core components whose functions may be representative functions of the respective module. Further, GO enrichment terms for “biological processes” are provided with the fold-enrichment and FDR values to infer the overall function of the co-expressed functional cluster.

#### By Conditions

To explore important stress-responsive genes, the user can browse NetREx “Condition-wise”. On selecting the tissue and stress/hormone treatment, the user can fetch the list of DEGs (in tabular format) for the corresponding tissue and condition (**Table 3**). Tables containing gene information and link to IC4R expression database, gene function (GO, MapMan and KEGG) and fold-change along with *p*-value across time-points are provided. The fold-change tables can be sorted by fold-change or *p*-value to identify most significant up- or down-regulated genes for the chosen condition at different time-points.

#### By Pathways

This is one of the attractive features of NetREx by which hierarchical KEGG pathways can be explored. After selecting the appropriate tissue and condition, the user may select a certain pathway of interest. Genes of the selected pathway that are DEGs for at least two time-points (for chosen stress/treatment condition) can be filtered for further network analysis.

## 4. Discussion

### Querying NetREx - A Case Study

To illustrate the efficacy of NetREx, we consider a set of drought-responsive genes in rice. The selected genes belong to the ABA signalosome complex comprising of PYR/PYL receptors, PP2Cs, SnRK2 kinases and ABF/bZIP transcription factors, obtained from the KEGG database (pathway ID: dosa04075) (39,40) for this case study. The ABA signal transduction pathway is one of the key mechanisms by which plants respond to environmental stresses like drought. Several studies indicate that the central signaling module comprises three protein classes: Pyracbactin Resistance/Pyracbactin resistance-like/Regulatory Component of ABA Receptor (PYR/PYL/RCARs) proposed to be the ABA receptors and the regulatory proteins, *viz*., Protein Phosphatase 2Cs (PP2Cs) which act as negative regulators together with SNF1-related protein kinases 2 (SnRKs) which are positive regulators (**Figure 2**). Increase in ABA levels during stress leads to the PYR/PYL/RCAR-PP2C complex formation causing inhibition of PP2C activity, thereby allowing activation of SnRKs which target the functional proteins like membrane proteins, ion channels and transcription factors, and facilitate transcription of ABA-responsive genes. Thus, network-based resources such as NetREx can enable us to query the coordinated interactions of regulatory genes and their functional targets which further trigger downstream processes.

**Figure 2.**
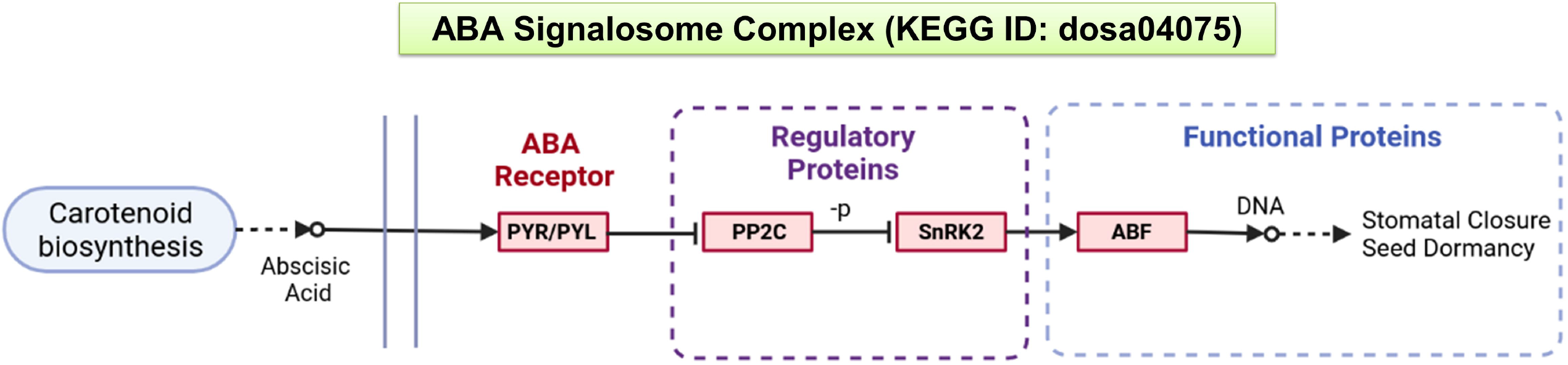
A schematic representation of the ABA signalling pathway from KEGG database

A total of 41 rice genes from KEGG (pathway ID: dosa04075) are queried in NetREx in “root” and “shoot” tissues under “drought” stress. Of these 13 and 17 genes respectively mapped to the root and shoot networks (**Table 6)**. The filtered gene sets (13 and 17) are a union of DEGs across all time points for the chosen condition (drought) and tissue (root and shoot). The “invalid genes” on the other hand either have very low expression values or are not differentially expressed in at least two time-points and hence not considered in the network construction in NetREx (**Table 2**).

**Table 6:** Shoot- and root-specific subnetworks of ABA Signalosome Complex extracted using NetREx under drought stress.

| RAP-DB Id | Gene Name | Component of the ABA Signalosome | Degree |
| --- | --- | --- | --- |
| **Shoot Network** |  |  |  |
| Os09g0325700 | OsSIPP2C1 | PP2C phosphatase | 7 |
| Os01g0583100 | OsPP2C06 | PP2C phosphatase | 6 |
| Os03g0268600 | OsPP2C30 | PP2C phosphatase | 6 |
| Os05g0537400 | OsPP2C50 | PP2C phosphatase | 6 |
| Os01g0656200 | OsPP2C08 | PP2C phosphatase | 5 |
| Os01g0869900 | SAPK4 | SnRK2 protein kinase | 5 |
| Os02g0766700 | OsbZIP23 | bZIP TF | 5 |
| Os06g0211200 | OsAREB1 | bZIP TF | 2 |
| Os07g0622000 | SAPK2 | SnRK2 protein kinase | 1 |
| Os05g0213500 | OsPYL5 | ABA receptor | 1 |
| Os01g0867300 | OsbZIP10 | bZIP TF | 1 |
| Os02g0551100 | SAPK6 | SnRK2 protein kinase | 1 |
| Os01g0813100 | HBF2 | bZIP TF | 0 |
| Os10g0564500 | SAPK3 | SnRK2 protein kinase | 0 |
| Os03g0390200 | SAPK1 | SnRK2 protein kinase | 0 |
| Os05g0437700 | OsbZIP40 | bZIP TF | 0 |
| Os02g0255500 | OsPYL3 | ABA receptor | 0 |
| **Root Network** |  |  |  |
| Os01g0656200 | OsPP2C08 | PP2C phosphatase | 7 |
| Os09g0325700 | OsSIPP2C1 | PP2C phosphatase | 7 |
| Os05g0537400 | OsPP2C50 | PP2C phosphatase | 7 |
| Os03g0268600 | OsPP2C30 | PP2C phosphatase | 7 |
| **Os01g0583100** | OsPP2C06 | PP2C phosphatase | 5 |
| **Os01g0867300** | OsbZIP10 | bZIP TF | 5 |
| **Os02g0766700** | OsbZIP23 | bZIP TF | 5 |
| **Os06g0211200** | OsAREB1 | bZIP TF | 5 |
| **Os02g0255500** | OsPYL3 | ABA receptor | 0 |
| **Os02g0551100** | SAPK6 | SnRK2 protein kinase | 0 |
| **Os07g0622000** | SAPK2 | SnRK2 protein kinase | 0 |
| **Os10g0573400** | OsPYL1 | ABA receptor | 0 |
| **Os09g0456200** | OsbZIP72 | bZIP TF | 0 |

In the **Expression Viewer**, fold-change of the filtered (valid) genes is displayed as heatmaps, shown in **Figure 3**. For the root tissue (**Figure 3 (A))**, it is observed that majority of the DEGs are strongly up-regulated as early as 1 hr time-point. However, at 3 hr, a decrease in fold-change is observed which is probably due to transcriptomic and metabolic reprogramming. For the shoot tissue (**Figure 3 (B))** it is observed that most genes are not induced at 1hr time-point but gradually the fold-change increases at later time-points. This indicates that response to drought stress is induced in root tissue earlier compared to shoot tissue.

**Figure 3.**
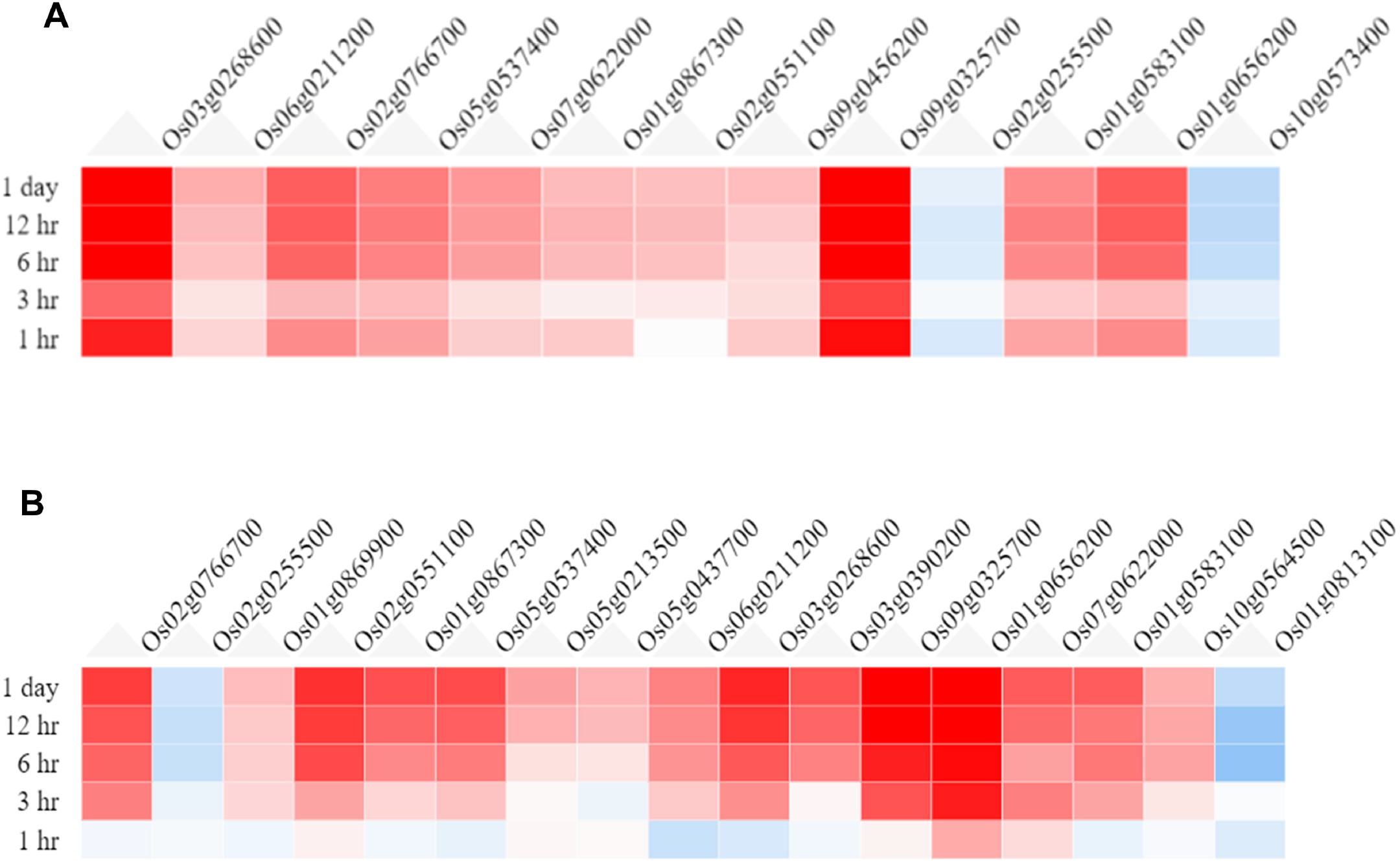
Heatmaps from Expression Viewer for (A) Root tissue and (B) Shoot tissue under Drought stress. The figure depicts transcriptional changes across time-points after drought stress with genes in red having high positive fold-change and those in blue having high negative fold-change with respect to control (0h, no stress)

Using “**Network Viewer**” module in NetREx, user can observe stress and tissue-specific view of the HRR-100 network. The expression status of the queried genes can be viewed in a time-specific manner by choosing the time-points 1h (default), 3h, 6h, 12h and 1day. In **Figure 4**, the “default view” indicates that among the 13 and 17 DEGs that mapped to the drought HRR-100 network in root (**Figure 4 (A))** and shoot (**Figure 4 (B))** respectively, 8 genes are seen to form the largest connected component in both these networks. In **Figure 5** is shown the connectivity information between genes, using the colour coding scheme: up-regulated (in red) and down-regulated (in blue) genes at 1h, 3h and 1day. For example, in the shoot network, the high-degree gene OsSIPP2C1 is induced at 3h of stress, while it is induced early on (at 1h) in root network. Interestingly, it was observed in previous studies that OsSIPP2C1 is negatively regulated by ABL1 which is involved in abiotic stress and panicle development in rice (41). In the root network, the components are more tightly connected with the PP2Cs *viz*., OsSIPP2C1, OsPP2C50, OsPP2C30 and OsPP2C08 with the highest degree genes interacting with the TFs OsAREB1, OsbZIP23 and OsABF1. Also, most of the genes are up-regulated at 1h time-point (**Figure 5**) and are probably early response genes in root tissue, except OsAREB1 which seems to be a late response gene activated only at 6h. For the shoot network, the response is slower as discussed above and the SNRK2 protein kinase, SAPK4, is not induced until 12h. While majority of the valid genes are up-regulated in shoot and root, a few genes are observed to be down-regulated. These include OsPYL3 (Os02g0255500), down-regulated both in root and shoot tissues. Indeed, this gene has been shown to be down-regulated in drought-susceptible rice genotype under abiotic stress conditions, while over-expression of this gene in Arabidopsis led to improved tolerance in cold and drought stress conditions (42). The shoot-specific TF HBF2 is also down-regulated in the shoot network (6h, not shown in the figure), while the root-specific OsPYL1 gene is down-regulated at 1h in the root network (Fig 2C). Among the 13 root and 17 shoot, 11 genes are common and differentially expressed in the two tissues. Interestingly, all the five 2C protein phosphatase (PP2C) proteins are common to both root and shoot, indicating their ubiquitous role of negative regulation of ABA (via SnRK2s and PYR/PYL/RCARs) in both the tissues (43). However, in terms of network-concepts, connectivity between the common gene sets differ between the two tissues as observed in **Figure 4**. Among the six shoot-specific DEGs, the bZIP TF HBF2 has been shown to be highly expressed in shoot apical meristem (44). On the other hand, the root-specific gene OsPYL1 was shown to interact with OsABIL2 that has a role in root development (42).

**Figure 4.**
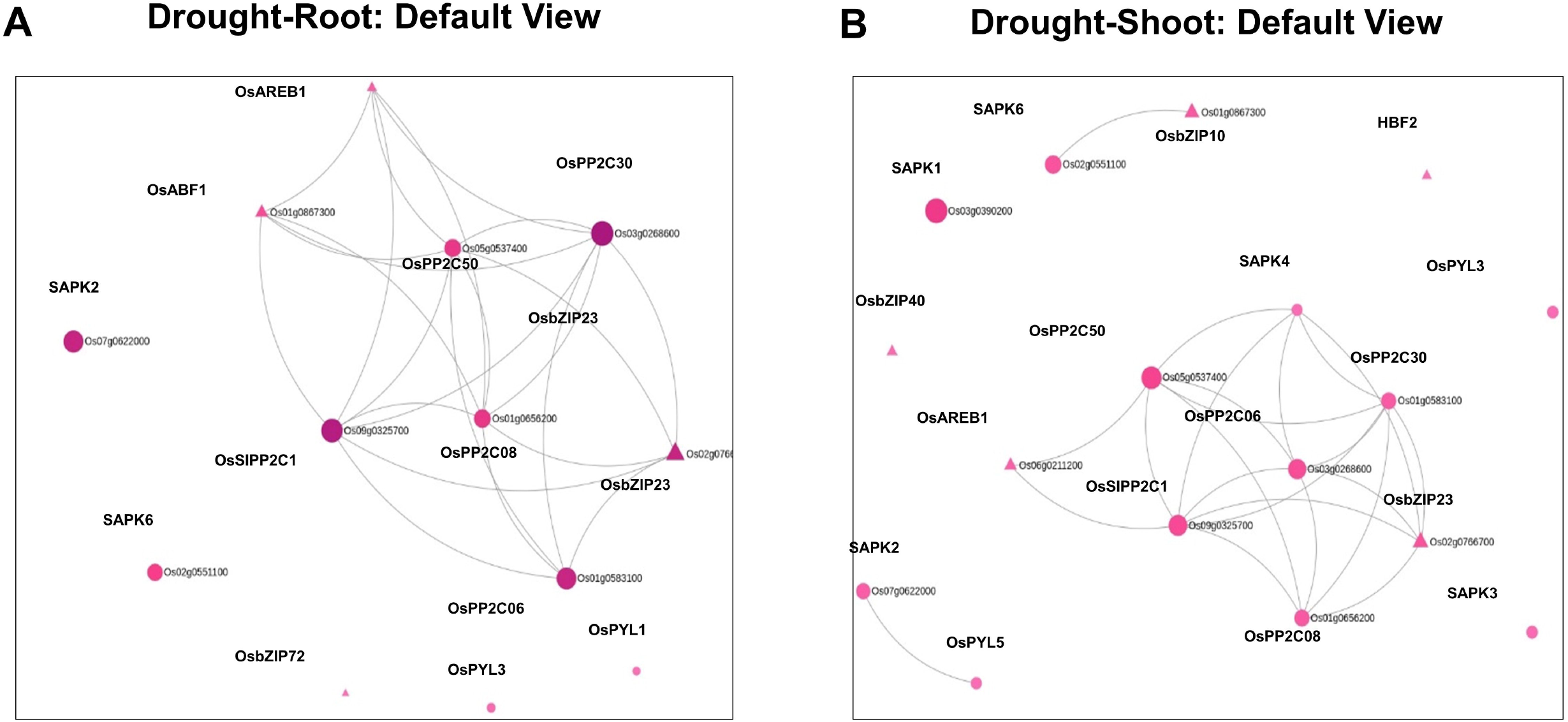
Default views for Network Viewer from (A) Root tissue and (B) Shoot tissue under Drought stress.

**Figure 5.**
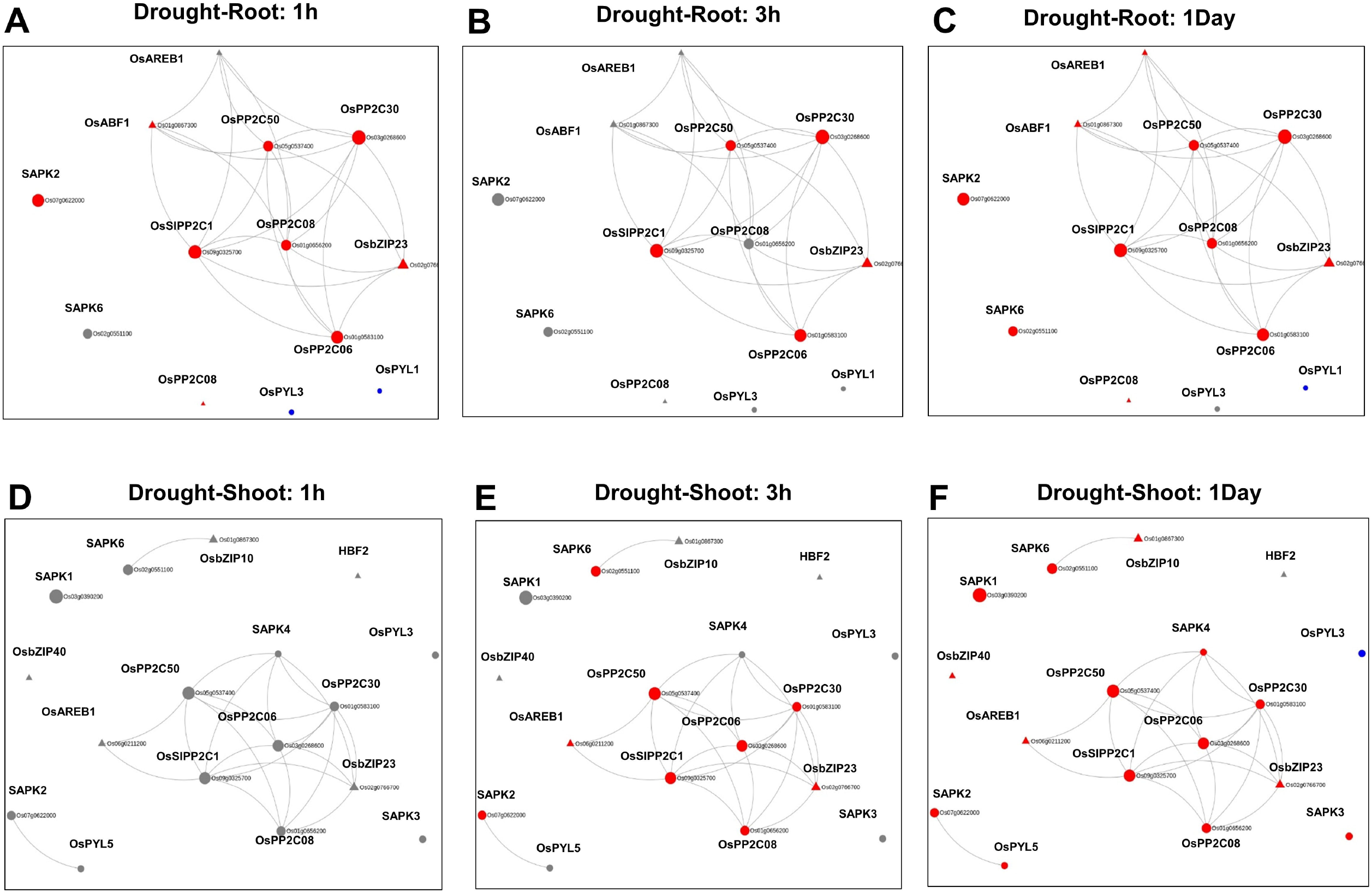
Time-point specific views from Network Viewer for Root and Shoot tissues at 1h, 3h and 1Day. The figure provides a comparative view of the transcriptitonal changes for the different time-points along with tissue-specific gene connectivities

Clusters of co-expressed genes often represent coordinated biological processes. Analysis of gene co-expression networks have been helpful in functional annotation of uncharacterized genes (36,45), prioritization of candidate genes (46,47), inferring biological processes, e.g., metabolic pathways (48,49), stress response mechanisms (50,51), cell wall metabolism (52,53), etc. For example, out of the 13 genes in the root network, 8 genes belong to the root-specific Magenta module, shown in **Figure 6**. Incidentally, they also form the largest component in the network (**Figure 6 (A))**. The remaining five genes with zero degree belong to GreenYellow, Yellow (2 genes), Blue and the Brown module. To obtain further details on the root-specific Magenta module, we use the “Browse” option in NetREx for “Root” tissue. Some of the significant GO terms include “regulation of transcription, DNA-templated” (GO:0006355, FDR= 8.56e-04) and “abscisic acid-activated signaling pathway” (GO:0009738, FDR= 7.64e-03) indicating the relevance of Magenta module in drought stress. Similarly, for the shoot network, out of 17 genes, 8 belong to the shoot-specific Turquoise and 4 to the shoot-specific Salmon modules. The Salmon module harbors genes involved in Dephosphorylation (GO:0016311, FDR= 1.98e-02), while the Turquoise module is involved in a number of stress-responsive processes including ER-associated misfolded protein catabolic process (GO:0071712, FDR= 8.83E-03), regulation of response to stress (GO:0080134, FDR=3.51e-05), etc. The Module Viewer allows further exploration of genes belonging to the respective modules. For example, the above analysis indicates that genes of root-specific Magenta module maybe biologically relevant for drought stress. On exploring the top 100 highly connected genes of the module, we see several known TF family genes like bZIPs and HSFs are important hubs of this module. Along with these, several “Conserved hypothetical proteins’ lacking detailed functional annotations are also part of the hubs. The associations of these genes with known TFs can be further queried in NetRex along with their expression profiles across different conditions and different tissues (IC4R Expressions).

**Figure 6.**
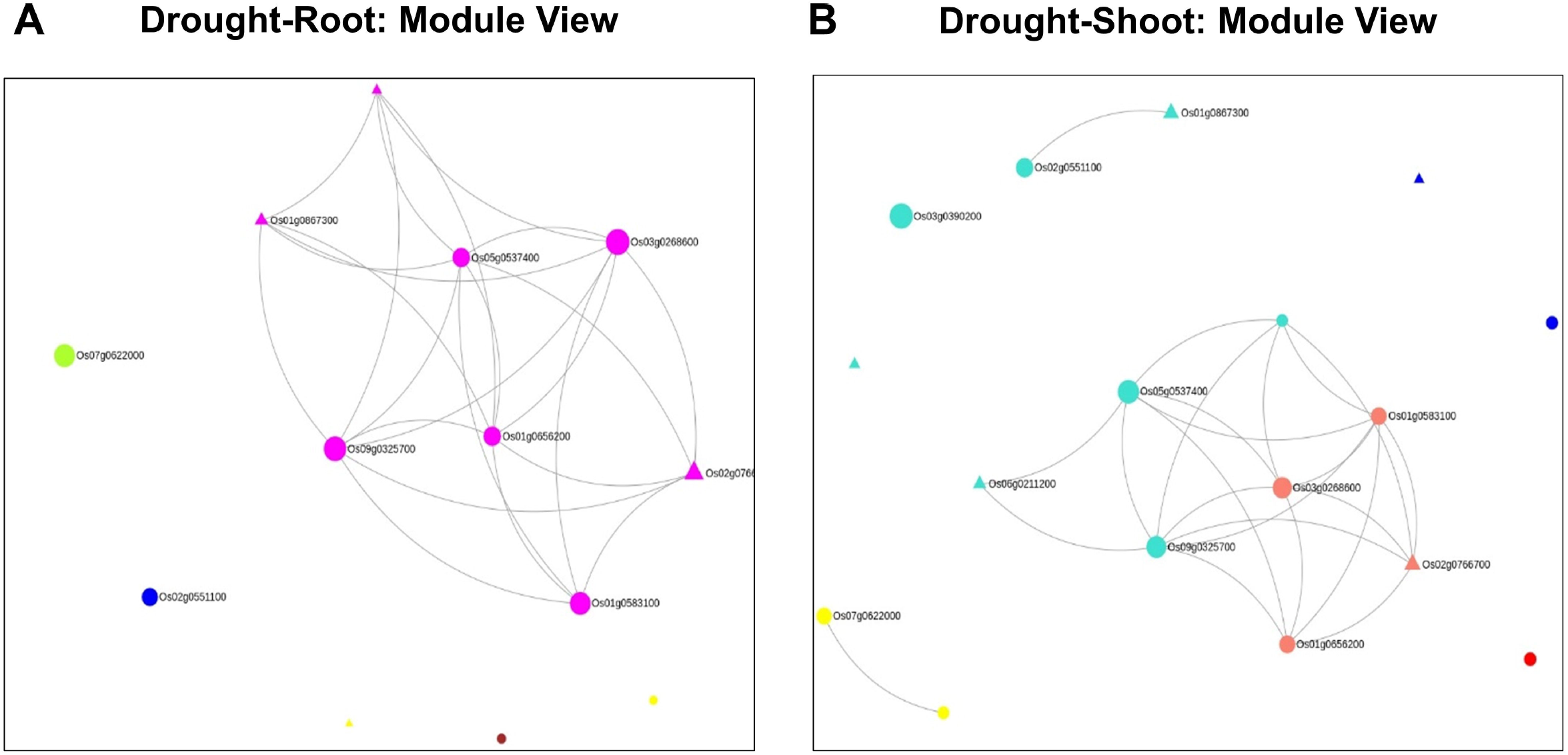
Module views of the Network Viewer for (A) Root tissue and (B) Shoot tissue under Drought stress.

In systems biology, network neighbourhood analysis is an important aspect as it facilitates a “guilt-by-association” strategy by which we can find interesting genes which are closely interacting/co-expressed with the initial “seed genes”. In the **Neighborhood View** on the right panel, top 100 neighbours based on k_Total_, the connectivity in the whole network, of the 13 root-specific seed genes are fetched. The “seed genes” are indicated with green borders in **Figure 7 (A)**. To infer the overall function of the subnetwork (13 query genes and their respective 100 neighbours), we performed GO enrichment analysis. As expected, positive regulation of abscisic acid-activated signalling pathway (GO:0009789, FDR=9.50E-03) was the most enriched term. Two major clusters are clearly discernible (**Figure 7 (B)**) in this subnetwork. The first set consists of neighbourhood genes that majorly belong to the Magenta module (16) and these genes are in fact induced as early as 1h of drought stress (**Figure 8 (A))**. As indicated above, the Magenta module is involved in the ABA signalling and stress responsive pathways. The other cluster consists of mostly genes belonging to Blue and Red modules. These sets of genes are late response genes are induced as a result of downstream cellular and metabolic adjustments after the signalling components have been induced in the early time-points (**Figure 2 and Figure 8 (B)**). The Blue module includes stress-responsive genes that aid in autophagy of damaged proteins and cellular organelles (GO:0044805, FDR=6.74E-03) (54). The Red module harbours genes involved in the methylerythritol 4-phosphate (MEP) pathway of isoprenoid biosynthetic process leading to the production of carotenoids and various other secondary metabolites (GO:0019288, FDR= 1.32E-03).

**Figure 7.**
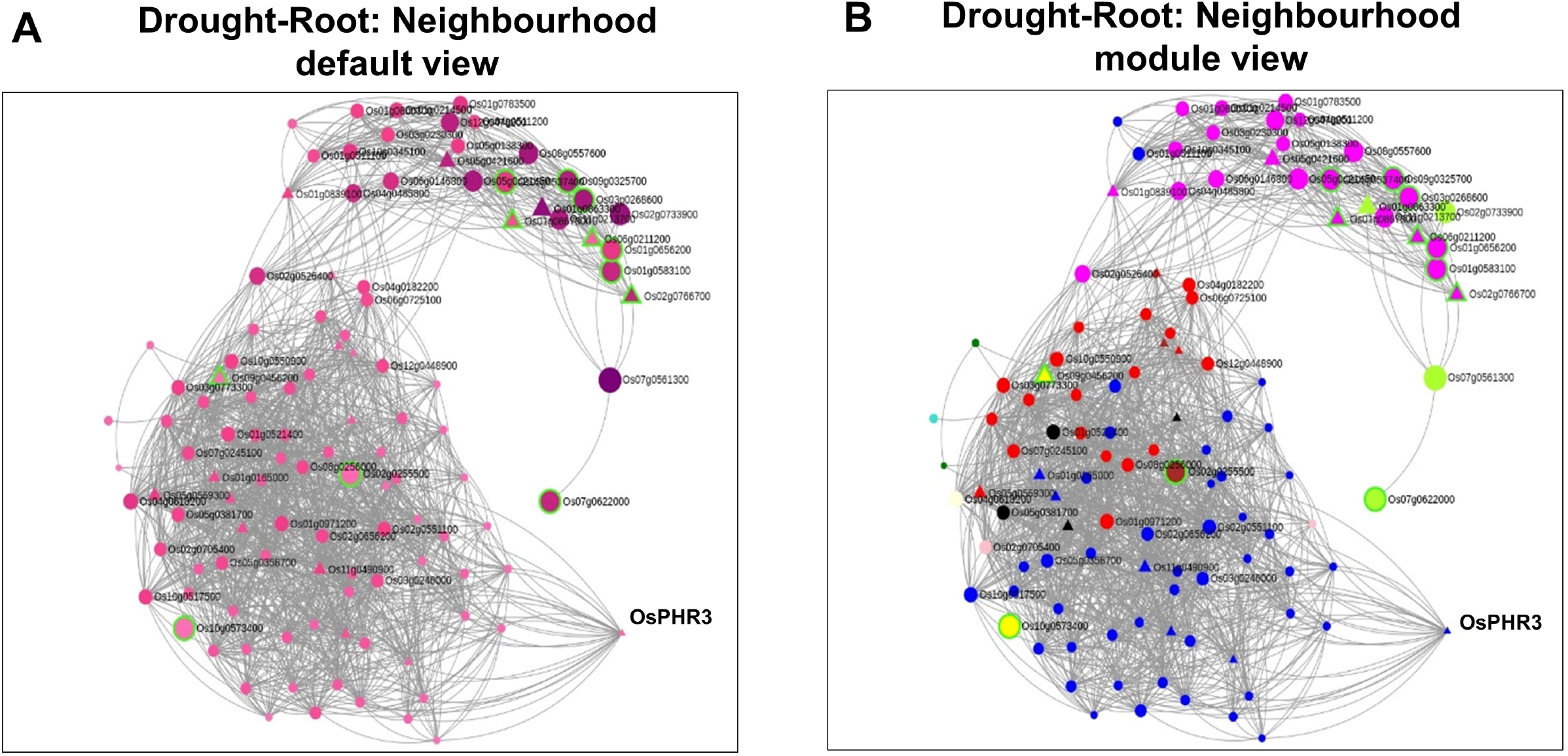
Neighborhood View of 13 root-specific genes with (A) default view and (B) Module views under

**Figure 8.**
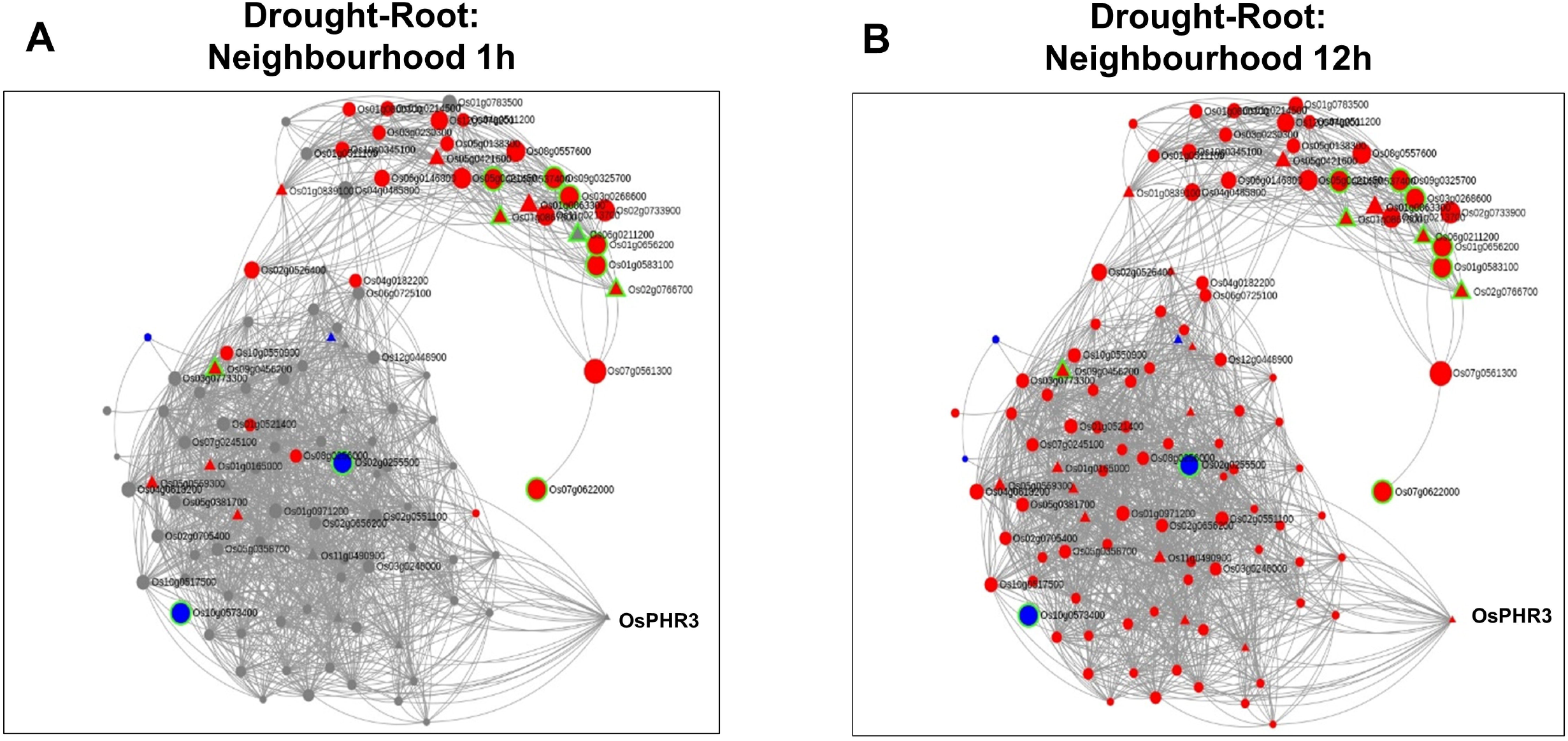
Time-point specific views for the extended neighbourhood for root-specific genes at (A) early timepoint of 1h and (B) late time-point of 12h drought stress

As discussed above, network neighbourhood helps us explore novel candidate genes that are absent in the initial query gene set. For example, the gene with the highest k_Total_ (weighted connectivity of a gene in the whole root network) is the transcription factor OsPHR3 (Os02g0139000) implicated in low Pi stress and in regulating Nitrogen homeostasis (55). To explore the transcriptomic dynamics of this gene in other stress conditions, we queried in NetREx again. Using the option to check the expression profile in ‘other conditions’ provided on the right panel in “Network Viewer”, we observed that this gene is up-regulated in osmotic stress (3-6h), flood stress (1h), ABA (1h to 1 day) and JA (1 and 3h) while it was also down-regulated in osmotic stress at 12h, flood stress (3-6h) and JA (6h to 1 day). Additionally, to explore the expression of this TF across rice growth stages and tissues, we used the IC4R link in the “Nodes Description” table. From **Figure 9** it may be noted that OsPHR3 exhibits higher expression in root and leaf tissues. On scanning the upstream 1kb of this gene using the PlantPan v2.0 database (56), we observed several bZIP binding motifs especially in the 500 kb upstream region **(Supplementary file1 (A))**. Further, concurrent with this, several binding sites for WRKY TFs were also detected in and around the same region, indicating that this TF may also be a target of biotic stress signaling cascades (**Supplementary file1 (B) and Supp. Table 8**). The network neighbourhood view of OsPHR3 was next explored. All its neighbours are observed to be up-regulated at 6h in root tissue under drought stress and all of them belong to the Blue module (**Supplementary file1 (A and B)**. Moreover, the top 10 neighbours are involved in functions like transferring phosphorus-containing groups, carbon-nitrogen ligase activity as well as drought and biotic stress (**Supp. Table 8)**. Literature survey revealed that this TF has not been functionally characterized in multiple stress conditions such as drought and warrants further investigation.

**Figure 9.**
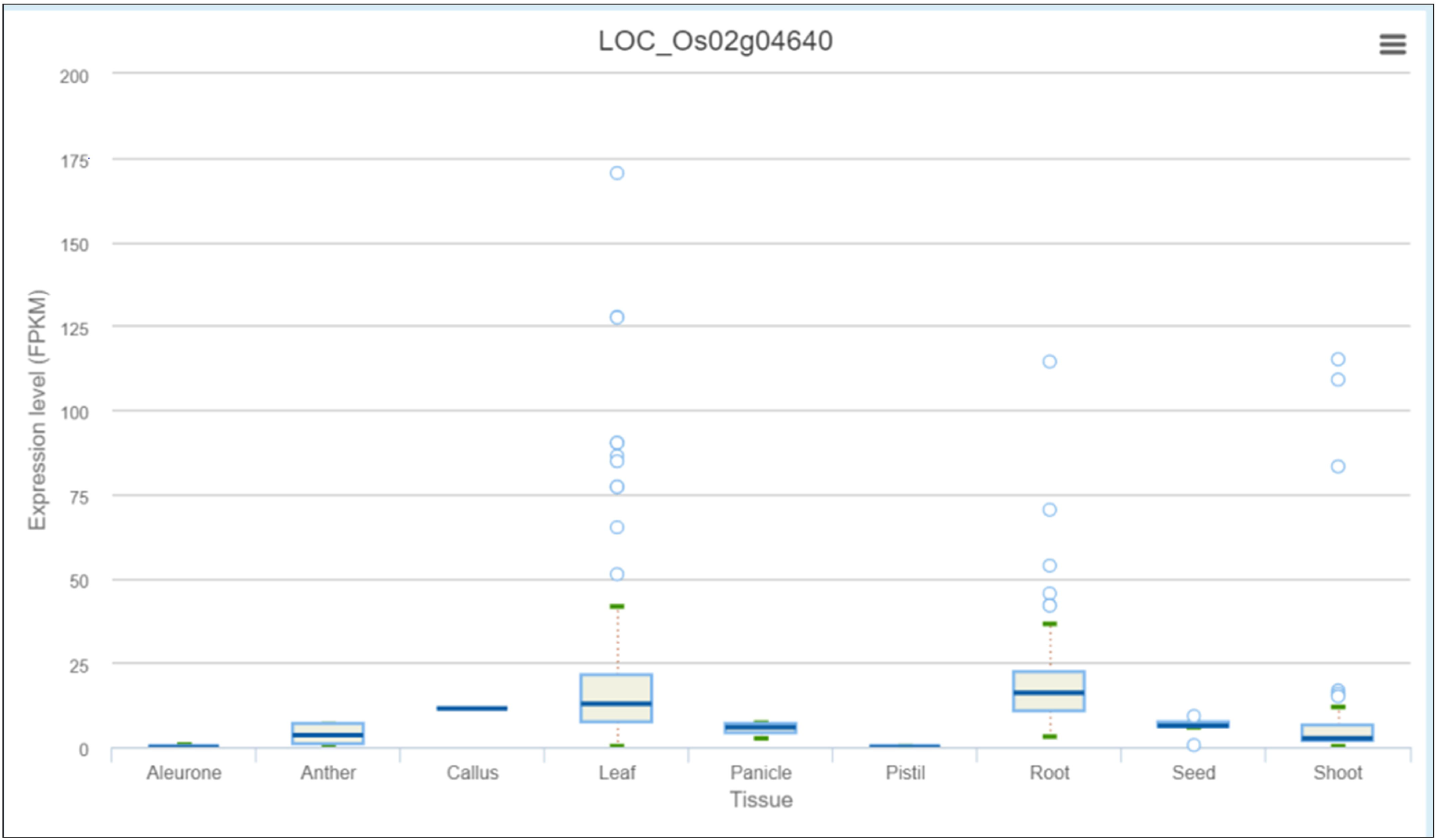
Expression profile of OsPHR3 from IC4R across different tissues and development stages.

A similar analysis was carried out with 17 shoot-specific DEGs and the Network View is shown in **Figure 10 (A)**. It may be noted from the module view in **Figure 10 (B)** that majority of the genes belong to the Turquoise Module while a separate cluster is formed by genes of the Blue Module. Some of the most significant functions of the subnetwork include protein serine/threonine phosphatase activity (GO:1905183, FDR=6.58E-03), ABA signalling (GO:0009789, FDR=8.95E-03), cellular response to nitrogen starvation (GO:0006995, FDR=1.28E-03), etc. Here, the gene with the highest k_Total_ is OsAtg8 (Os07g0512200), which is a well characterized gene involved in autophagy and protein degradation (57). However, the role of this gene with respect to drought is yet to be explored. An important point to be noted is that we extracted extended root and shoot networks under drought stress using the same “seed” genes involved in ABA signalling. Needless to say, both the subnetworks had GO terms enriched for “abscisic acid-activated signaling pathway” (GO:0009738), “regulation of response to water deprivation” (GO:2000070), “cellular response to hormone stimulus” (GO:0032870) and so on. However, more specific tissue-specific GO terms like “photosynthesis and dark reaction” (GO:0019685, FDR= 3.15E-02), “gluconeogenesis” (GO:0006094, FDR= 3.60E-03), “cellular response to nitrogen levels” (GO:0043562, FDR= 4.21E-05), etc. for the shoot tissue and “cellular response to reactive oxygen species” (GO:0034614, FDR= 3.57E-02), “fatty acid oxidation” (GO:0019395, FDR= 5.41E-03), etc. for the root tissue were noted. Another major difference in the **Neighborhood View** of these genes in root and shoot tissues is the presence of down-regulated genes in the shoot network as compared to root and the gradual activation/repression of these genes in shoot (**Figure 11 (A to D))** in contrast to root (**Figure 8 (A** and **B)**. The down-regulated genes in shoot (21 genes) majorly belong to the **Blue** module (**Figure 10 (B**)) with at least 6 genes annotated to be involved in photosynthesis, indicating that this process is preferentially switched-off in the green tissues under drought stress. Further exploration of the **Blue** module of the shoot HRR network using the “Module Wise” page in NetREx revealed several interesting shoot-specific GO terms like “photosystem II repair” (GO:0010206, FDR= 9.62E-04), “photosystem II assembly” (GO:0010207, FDR= 4.41E-03), “regulation of photosynthesis, light reaction” (GO:0042548, FDR= 3.80e-03), etc. Adverse effects of abiotic stress conditions like drought on the photosynthetic machinery with harmful effects on the overall growth and yield of the crop is well documented (58,59). This confidently explains the functional differences of the root and shoot subnetworks from the extended ABA signalosome analysis discussed above. Moreover, the expression profiles of the genes across time-points and their corresponding tissue-specific network connectivities enable one to confidently explore the temporal and functional space of the genes and arrive at relevant conclusions.

**Figure 10.**
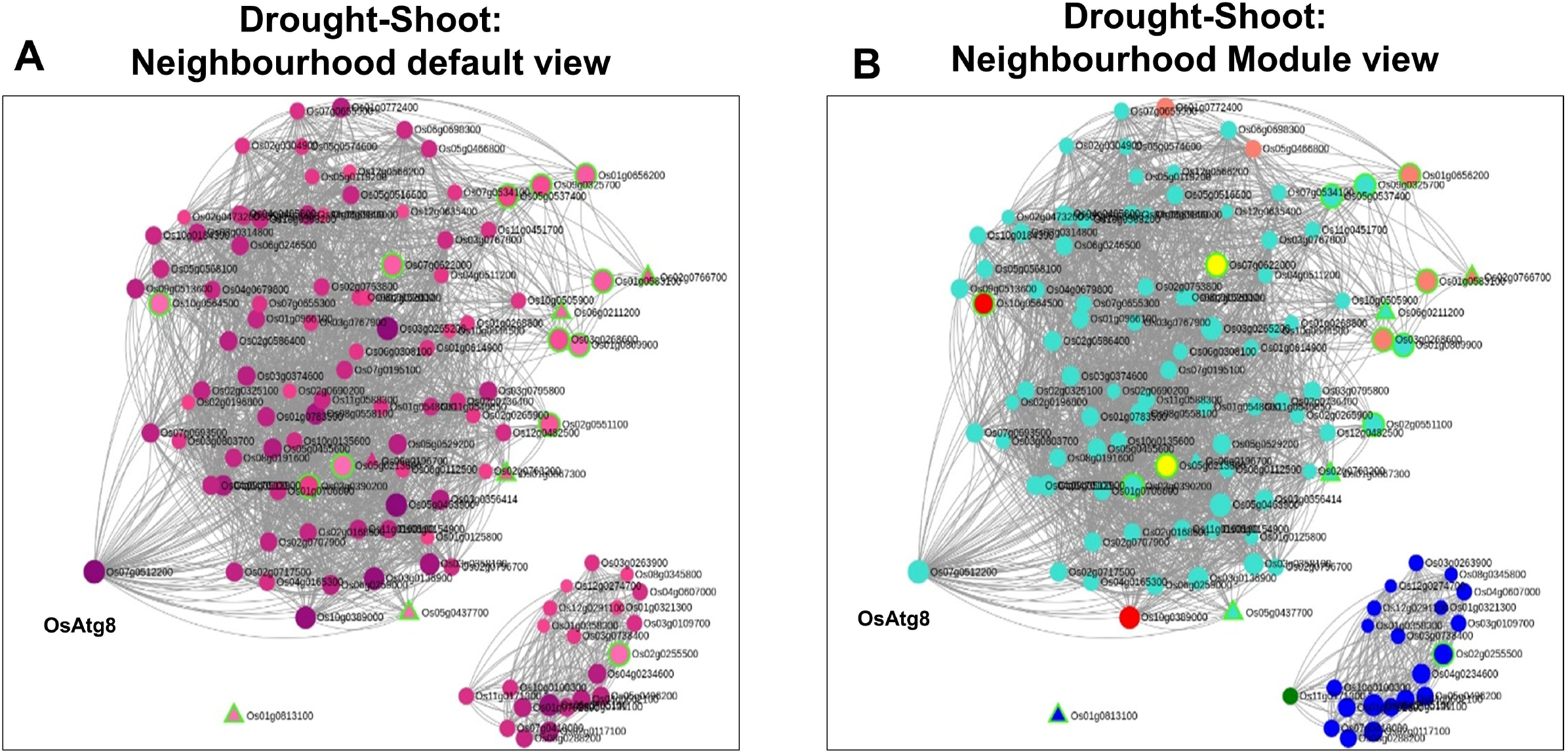
Neighborhood View of 17 shoot-specific genes with (A) default view and (B) Module views under Drought stress

**Figure 11.**
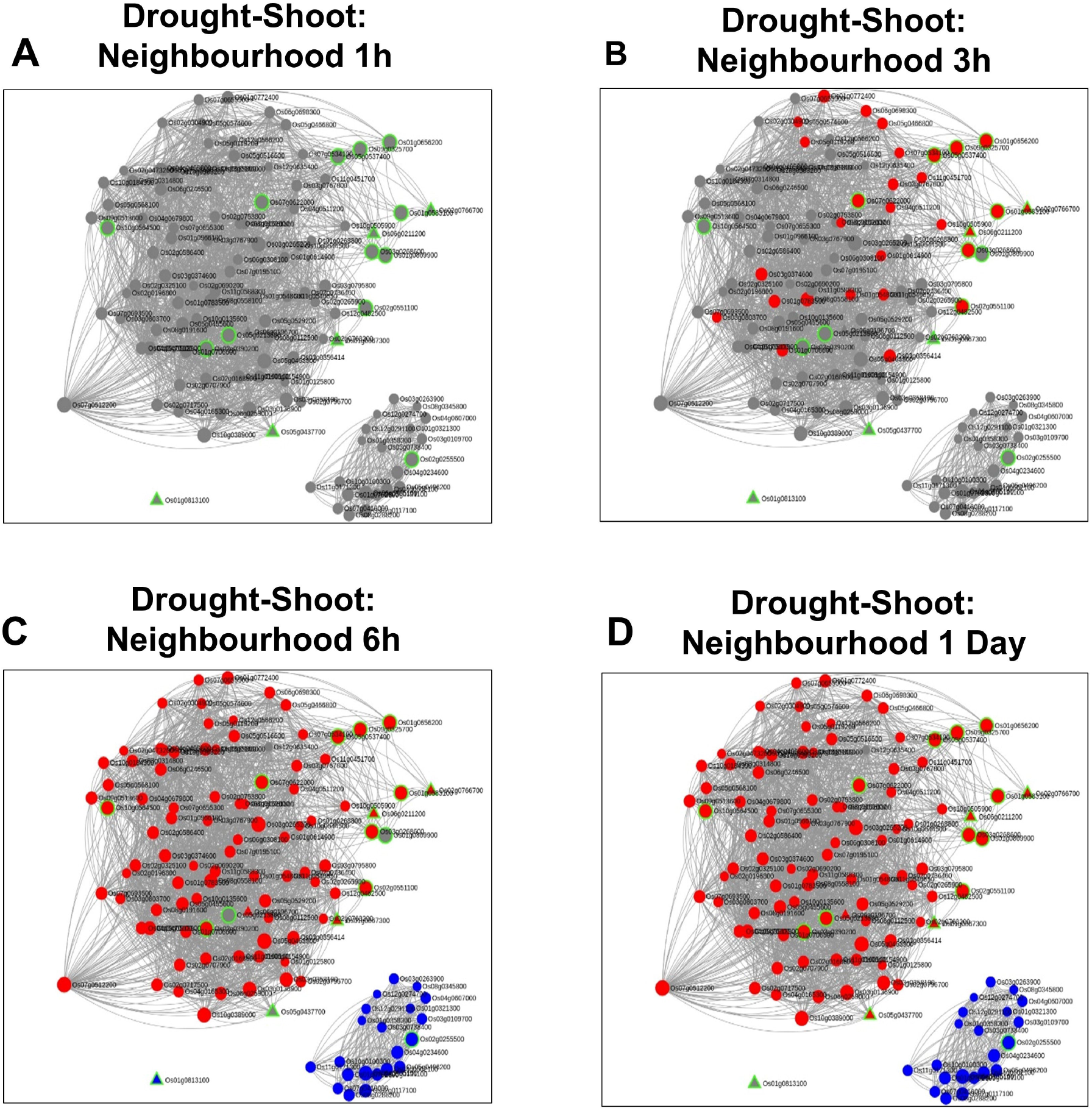
Time-point specific views for the extended neighbourhood for shoot-specific genes at early timepoints (1 and 3h) and late time-points (6h and 1day)

## Conclusion

NetREx is a freely accessible web-server for biologists to conveniently explore the global rank-based stress networks in a tissue-specific manner. The resource has been constructed using high-quality RNA-seq data from the TENOR database generated using homogeneous experimental protocols. In NetREx, substantial emphasis has been given to explore the networks through various perspectives such as exploring gene expression profiles (Expression Viewer heatmaps and Network Viewer in a time-point specific manner), network connectivity (Network Viewer and Neighbourhood Viewer), identification of novel stress-responsive candidates (Neighbourhood Viewer), functional analysis of genes (browsing NetREx by modules and pathways) and comparative analysis across stress conditions (supported in Network Viewer mode). The gene attributes displayed in the different modules have been extensively cross-linked to various other resources to provide additional information to the users. Our analysis indicates that the rank-based networks in NetREx are biologically relevant wherein the tissue and stress-specific information is effectively retained. Network-based subnetwork analysis and gene prioritization using NetREx will therefore be a significant resource to study complex phenotypes associated with stress-response in rice.

## Conflict of interest

The authors declare no conflict of interest.

## Notes

https://bioinf.iiit.ac.in/netrex/index.html

### Competing Interest Statement

The authors have declared no competing interest.

## References

1. Rabbani, M.A., Maruyama, K., Abe, H., et al. (2003) Monitoring expression profiles of rice genes under cold, drought, and high-salinity stresses and abscisic acid application using cDNA microarray and RNA gel-blot analyses. Plant physiology, 133, 1755–67.

2. Rensink, W.A., Iobst, S., Hart, A., et al. (2005) Gene expression profiling of potato responses to cold, heat, and salt stress. Functional & Integrative Genomics, 5, 201–207.

3. Achuo, E.A., Prinsen, E. and Hofte, M. (2006) Influence of drought, salt stress and abscisic acid on the resistance of tomato to Botrytis cinerea and Oidium neolycopersici. Plant Pathology, 55, 178–186.

4. Rasmussen, S., Barah, P., Suarez-Rodriguez, M.C., et al. (2013) Transcriptome responses to combinations of stresses in Arabidopsis. Plant physiology, 161, 1783–94.

5. Mantri, N.L., Ford, R., Coram, T.E., et al. (2007) Transcriptional profiling of chickpea genes differentially regulated in response to high-salinity, cold and drought. BMC Genomics, 8, 303.

6. Sato, Y., Takehisa, H., Kamatsuki, K., et al. (2013) RiceXPro Version 3.0: expanding the informatics resource for rice transcriptome. Nucleic Acids Research, 41, D1206–D1213.

7. Priya, P. and Jain, M. (2013) RiceSRTFDB: a database of rice transcription factors containing comprehensive expression, cis-regulatory element and mutant information to facilitate gene function analysis. Database, bat027.

8. Waese, J., Fan, J., Pasha, A., et al. (2017) ePlant: Visualizing and Exploring Multiple Levels of Data for Hypothesis Generation in Plant Biology. The Plant cell, 29, 1806–1821.

9. Xia, L., Zou, D., Sang, J., et al. (2017) Rice Expression Database (RED): An integrated RNA-Seq-derived gene expression database for rice. Journal of Genetics and Genomics, 44, 235–241.

10. Kawahara, Y., Oono, Y., Wakimoto, H., et al. (2016) TENOR: Database for Comprehensive mRNA-Seq Experiments in Rice. Plant and Cell Physiology, 57, e7–e7.

11. Serin, E.A.R., Nijveen, H., Hilhorst, H.W.M., et al. (2016) Learning from co-expression networks: Possibilities and challenges. Learning from co-expression networks: Possibilities and challenges. Frontiers in Plant Science, 7.

12. Mutwil, M., Usadel, B., Schütte, M., et al. (2010) Assembly of an interactive correlation network for the Arabidopsis genome using a novel heuristic clustering algorithm. Plant physiology, 152, 29–43.

13. Obayashi, T. and Kinoshita, K. (2009) Rank of Correlation Coefficient as a Comparable Measure for Biological Significance of Gene Coexpression. DNA Research, 16, 249–260.

14. Liesecke, F., Daudu, D., Dugé de Bernonville, R., et al. (2018) Ranking genome-wide correlation measurements improves microarray and RNA-seq based global and targeted co-expression networks. Scientific Reports, 8, 10885.

15. Mutwil, M., Øbro, J., Willats, W.G.T., et al. (2008) GeneCAT—novel webtools that combine BLAST and co-expression analyses. Nucleic Acids Research, 36, W320–W326.

16. Obayashi, T., Aoki, Y., Tadaka, S., et al. (2018) ATTED-II in 2018: A Plant Coexpression Database Based on Investigation of the Statistical Property of the Mutual Rank Index. Plant and Cell Physiology, 59, e3–e3.

17. Proost, S. and Mutwil, M. (2018) CoNekT: an open-source framework for comparative genomic and transcriptomic network analyses. Nucleic Acids Research, 46, W133–W140.

18. Proost, S. and Mutwil, M. (2017) PlaNet: Comparative Co-Expression Network Analyses for Plants. Methods in molecular biology (Clifton, N.J.), 1533, 213–227.

19. Wong, D.C.J., Sweetman, C., Drew, D.P., et al. (2013) VTCdb: a gene co-expression database for the crop species Vitis vinifera (grapevine). BMC genomics, 14, 882.

20. Lee, T., Yang, S., Kim, E., et al. (2015) AraNet v2: An improved database of co-functional gene networks for the study of Arabidopsis thaliana and 27 other nonmodel plant species. Nucleic Acids Research, 43, D996–D1002.

21. Lee, T., Oh, T., Yang, S., et al. (2015) RiceNet v2: An improved network prioritization server for rice genes. Nucleic Acids Research, 43, W122–W127.

22. Lee, T., Lee, S., Yang, S., et al. (2019) MaizeNet: a co-functional network for network-assisted systems genetics in Zea mays. Plant Journal.

23. Obayashi, T., Nishida, K., Kasahara, K., et al. (2011) ATTED-II updates: condition-specific gene coexpression to extend coexpression analyses and applications to a broad range of flowering plants. Plant & cell physiology, 52, 213–9.

24. Fukushima, A., Nishizawa, T., Hayakumo, M., et al. (2012) Exploring tomato gene functions based on coexpression modules using graph clustering and differential coexpression approaches. Plant physiology, 158, 1487–502.

25. Martin, M. (2011) Cutadapt removes adapter sequences from high-throughput sequencing reads. EMBnetjournal, 17(1).

26. Kim, D., Langmead, B. and Salzberg, S.L. (2015) HISAT: a fast spliced aligner with low memory requirements. Nature Methods, 12, 357–360.

27. Sakai, H., Lee, S.S., Tanaka, T., et al. (2013) Rice Annotation Project Database (RAP-DB): An Integrative and Interactive Database for Rice Genomics. Plant and Cell Physiology, 54, e6–e6.

28. Conesa, A., Madrigal, P., Tarazona, S., et al. (2016) A survey of best practices for RNA-seq data analysis. Genome biology, 17, 13.

29. Liao, Y., Smyth, G.K. and Shi, W. (2014) featureCounts: an efficient general purpose program for assigning sequence reads to genomic features. Bioinformatics, 30, 923–930.

30. Love, M.I., Huber, W. and Anders, S. (2014) Moderated estimation of fold change and dispersion for RNA-seq data with DESeq2. Genome Biology, 15.

31. Anders, S. and Huber, W. (2010) Differential expression analysis for sequence count data. Genome Biology, 11, R106.

32. Mason, M.J., Fan, G., Plath, K., et al. (2009) Signed weighted gene co-expression network analysis of transcriptional regulation in murine embryonic stem cells. BMC Genomics, 10, 327.

33. Langfelder, P., Zhang, B. and Horvath, S. (2008) Defining clusters from a hierarchical cluster tree: the Dynamic Tree Cut package for R. Bioinformatics, 24, 719–720.

34. Mi, H., Muruganujan, A., Ebert, D., et al. (2019) PANTHER version 14: More genomes, a new PANTHER GO-slim and improvements in enrichment analysis tools. Nucleic Acids Research, 47, D419–D426.

35. Fernandez, N.F., Gundersen, G.W., Rahman, A., et al. (2017) Clustergrammer, a web-based heatmap visualization and analysis tool for high-dimensional biological data. Scientific Data 2017 4:1, 4, 1–12.

36. Sircar, S. and Parekh, N. (2015) Functional characterization of drought-responsive modules and genes in Oryza sativa: a network-based approach. Frontiers in genetics, 6, 256.

37. Kanehisa, M., Sato, Y., Kawashima, M., et al. (2016) KEGG as a reference resource for gene and protein annotation. Nucleic Acids Research, 44, D457–D462.

38. Usadel, B., Poree, F., Nagel, A., et al. (2009) A guide to using MapMan to visualize and compare Omics data in plants: a case study in the crop species, Maize. Plant, Cell & Environment, 32, 1211–1229.

39. Kanehisa, M. and Goto, S. (2000) KEGG: kyoto encyclopedia of genes and genomes. Nucleic acids research, 28, 27–30.

40. Umezawa, T., Sugiyama, N., Mizoguchi, M., et al. (2009) Type 2C protein phosphatases directly regulate abscisic acid-activated protein kinases in Arabidopsis. Proceedings of the National Academy of Sciences of the United States of America, 106, 17588–17593.

41. Li, Y.S., Sun, H., Wang, Z.F., et al. (2013) A novel nuclear protein phosphatase 2C negatively regulated by ABL1 is involved in abiotic stress and panicle development in rice. Molecular Biotechnology, 54, 703–710.

42. Li, C., Shen, H., Wang, T., et al. (2015) ABA Regulates Subcellular Redistribution of OsABI-LIKE2, a Negative Regulator in ABA Signaling, to Control Root Architecture and Drought Resistance in Oryza sativa. Plant and Cell Physiology, 56, 2396–2408.

43. Fujii, H., Chinnusamy, V., Rodrigues, A., et al. (2009) In vitro reconstitution of an abscisic acid signalling pathway. Nature, 462, 660–664.

44. v, B., d, M., d, G., et al. (2017) Antagonistic Transcription Factor Complexes Modulate the Floral Transition in Rice. The Plant cell, 29, 2801–2816.

45. Chandran, A.K.N., Bhatnagar, N., Yoo, Y.-H., et al. (2018) Meta-expression analysis of unannotated genes in rice and approaches for network construction to suggest the probable roles. Plant Molecular Biology, 96, 17–34.

46. Zeng, L., Liu, X., Zhou, Z., et al. (2018) Identification of a G2-like transcription factor, OsPHL3, functions as a negative regulator of flowering in rice by co-expression and reverse genetic analysis. BMC Plant Biology, 18, 157.

47. Schaefer, R., Michno, J.-M., Jeffers, J., et al. (2018) Integrating co-expression networks with GWAS to prioritize causal genes in maize. The Plant Cell, 30(12), 2922–2942.

48. Kautsar, S.A., Suarez Duran, H.G., Blin, K., et al. (2017) plantiSMASH: automated identification, annotation and expression analysis of plant biosynthetic gene clusters. Nucleic Acids Research, 45, W55–W63.

49. Wisecaver, J.H., Borowsky, A.T., Tzin, V., et al. (2017) A Global Coexpression Network Approach for Connecting Genes to Specialized Metabolic Pathways in Plants. The Plant cell, 29, 944–959.

50. Nounjan, N., Chansongkrow, P., Charoensawan, V., et al. (2018) High Performance of Photosynthesis and Osmotic Adjustment Are Associated With Salt Tolerance Ability in Rice Carrying Drought Tolerance QTL: Physiological and Co-expression Network Analysis. Frontiers in plant science, 9, 1135.

51. Tan, M., Cheng, D., Yang, Y., et al. (2017) Co-expression network analysis of the transcriptomes of rice roots exposed to various cadmium stresses reveals universal cadmium-responsive genes. BMC Plant Biology, 17, 194.

52. Wang, S., Yin, Y., Ma, Q., et al. (2012) Genome-scale identification of cell-wall related genes in Arabidopsis based on co-expression network analysis. BMC plant biology, 12, 138.

53. Ferreira, S.S., Hotta, C.T., Poelking, V.G. de C., et al. (2016) Co-expression network analysis reveals transcription factors associated to cell wall biosynthesis in sugarcane. Plant Molecular Biology, 91, 15–35.

54. Su, W., Bao, Y., Yu, X., et al. (2020) Autophagy and its regulators in response to stress in plants. Autophagy and its regulators in response to stress in plants. International Journal of Molecular Sciences, 21, 1–12.

55. Sun, Y., Luo, W., Jain, A., et al. (2018) OsPHR3 affects the traits governing nitrogen homeostasis in rice. BMC Plant Biology, 18, 1–15.

56. Chow, C.-N., Zheng, H.-Q., Wu, N.-Y., et al. (2016) PlantPAN 2.0: an update of plant promoter analysis navigator for reconstructing transcriptional regulatory networks in plants. Nucleic Acids Research, 44.

57. Su, W., Ma, H., Liu, C., et al. (2006) Identification and characterization of two rice autophagy associated genes, OsAtg8 and OsAtg4. Molecular biology reports, 33, 273–278.

58. Sircar, S. and Parekh, N. (2019) Meta-analysis of drought-tolerant genotypes in Oryza sativa: A network-based approach. PLOS ONE, 14, e0216068.

59. Zhou, Y., Lam, H.M. and Zhang, J. (2007) Inhibition of photosynthesis and energy dissipation induced by water and high light stresses in rice. Journal of Experimental Botany, 58, 1207–1217.

