## Supplementary figures and images for "NetREx – Network-based Rice Expression Analysis Server for Abiotic Stress Conditions"

### Supplementary file1

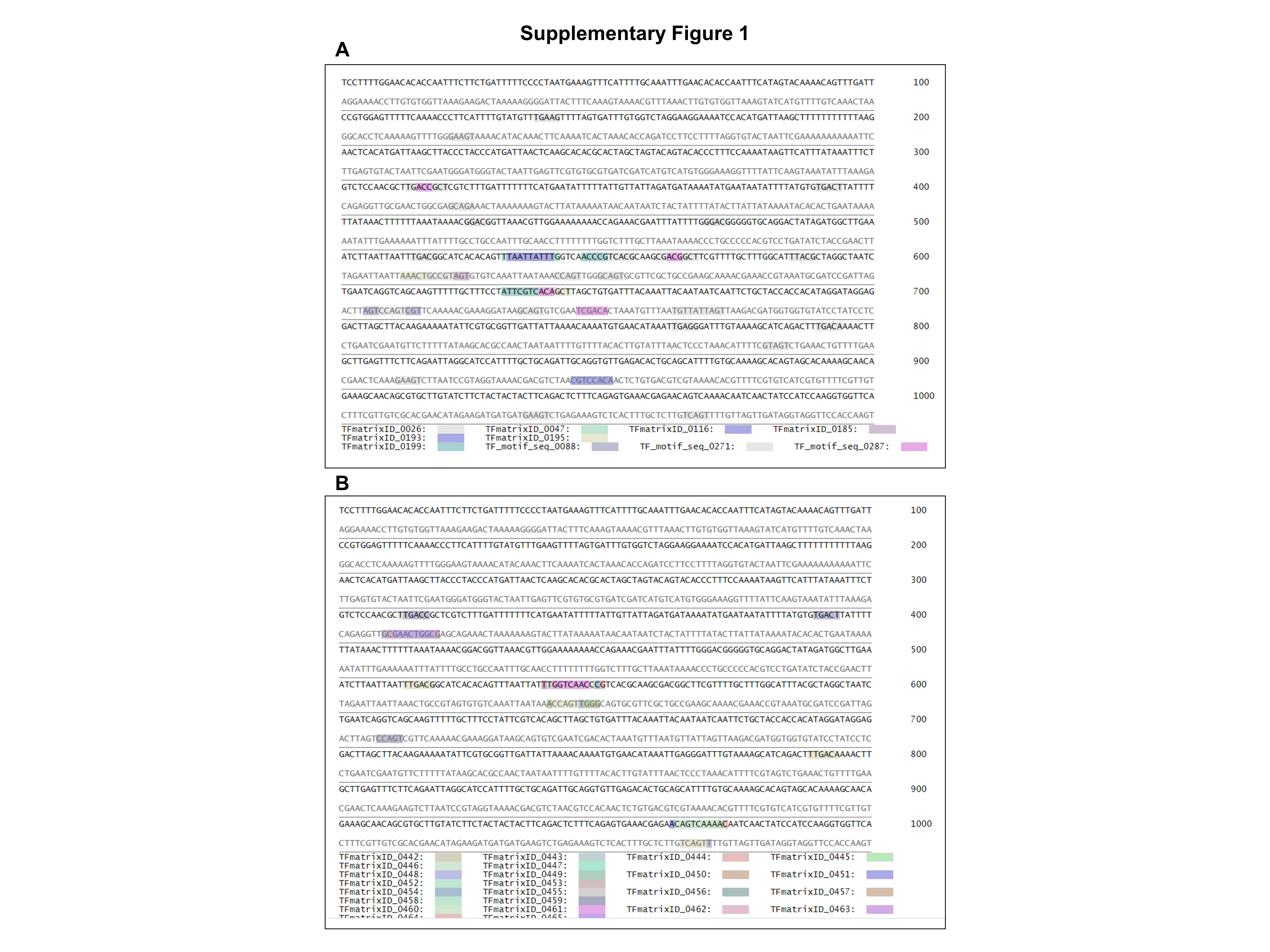

### Supplementary file2

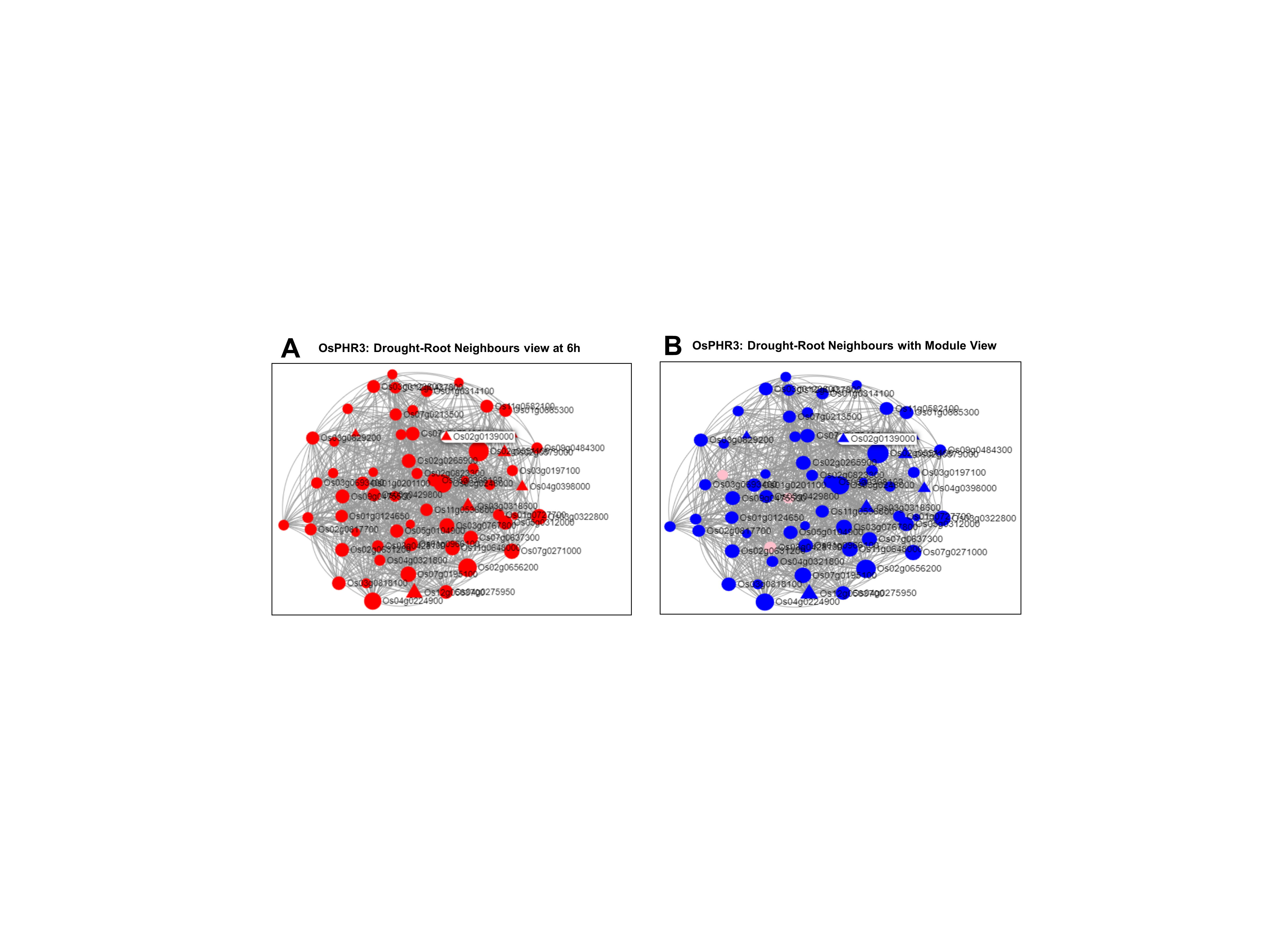
